# Development of a sensor for disulfide bond formation in diverse bacteria

**DOI:** 10.1101/2023.12.18.572236

**Authors:** Jocelyne Mendoza, Dyotima, Sally Abulaila, Cristina Landeta

**Author notes:** Correspondence should be addressed to Dr. Cristina Landeta.

## Abstract

In bacteria, disulfide bonds contribute to the folding and stability of proteins important for processes in the cellular envelope. In *E. coli*, disulfide bond formation is catalyzed by DsbA and DsbB enzymes. DsbA is a periplasmic protein that catalyzes disulfide bond formation in substrate proteins while DsbB is an inner membrane protein that transfers electrons from DsbA to quinones, thereby regenerating the DsbA active state. Actinobacteria including mycobacteria use an alternative enzyme named VKOR which performs the same function as DsbB. Disulfide bond formation enzymes, DsbA and DsbB/ VKOR represent novel drug targets because their inhibition could simultaneously affect the folding of several cell envelope proteins including virulence factors, proteins involved in outer membrane biogenesis, cell division, and antibiotic resistance. We have previously developed a cell-based and target-based assay to identify molecules that inhibit the DsbB and VKOR in pathogenic bacteria, using *Escherichia coli* cells expressing a periplasmic β-Galactosidase sensor (β-Gal^dbs^) which is only active when disulfide bond formation is inhibited. Here we report the construction of plasmids that allow fine-tuning of the expression of the β-Gal^dbs^ sensor and can be mobilized into other gram-negative organisms. As an example, when harbored in *P. aeruginosa* UCBPP-PA14, β-Gal^dbs^ behaves similarly as in *E. coli* and the biosensor responds to the inhibition of the two DsbB proteins. Thus, these β-Gal^dbs^ reporter plasmids provide a basis for identifying novel inhibitors of DsbA and DsbB/VKOR against multi-drug resistant, gram-negative pathogens and to further study oxidative protein folding in diverse gram-negative bacteria.

**Importance:** Disulfide bonds contribute to the folding and stability of proteins in the bacterial cell envelope. Disulfide bond-forming enzymes represent new drug targets against multidrug-resistant bacteria since inactivation of this process would simultaneously affect several proteins in the cell envelope, including virulence factors, toxins, proteins involved in outer membrane biogenesis, cell division, and antibiotic resistance. Identifying the enzymes involved in disulfide bond formation in gram-negative pathogens as well as their inhibitors can contribute to the much-needed antibacterial innovation. In this work, we developed sensors of disulfide bond formation for gram-negative bacteria. These tools will enable the study of disulfide bond formation and the identification of inhibitors for this crucial process in diverse gram-negative pathogens.

## Introduction

More than three decades ago the enzymes that introduce disulfide bonds were discovered thanks to a protein fusion that was developed as a complementary approach to studying membrane protein topology (1, 2). Froshauer S. *et al*., generated hybrid proteins of the membrane protein MalF with β-Galactosidase (β-Gal, LacZ). Cytoplasmic domain fusions exhibited high levels of β-Gal activity, whereas the periplasmic domain fusions expressed low activity. This approach complemented the use of PhoA fusions, which behaved oppositely to β-Gal and strengthened the evidence of membrane topology (1, 3, 4). One fusion, MalF-LacZ102 (aka β-Gal^dbs^ for disulfide bond-sensitive β-Gal) displayed low levels of β-Gal and it was found to be an in-frame hybrid of 198 amino acids of MalF attached by a three-amino acid linker (Gly-Asp-Pro) to the residue 8 of β-Gal (Figure 1) (1). The fusion occurred at the third transmembrane segment in the middle of the long hydrophilic domain thus, placing β-Gal in the periplasm (1, 5). When β-Gal^dbs^ is expressed in the periplasm of cells with an active disulfide bond formation system, β-Gal^dbs^ is inactivated with disulfide bonds, while an inactive disulfide bond machinery would produce active β-Gal^dbs^ (Figure 1). This fusion has been a crucial tool for the seminal discovery of the disulfide bond formation pathway in *Escherichia coli,* through two decades later to identify inhibitors of this process.

**Figure 1.**
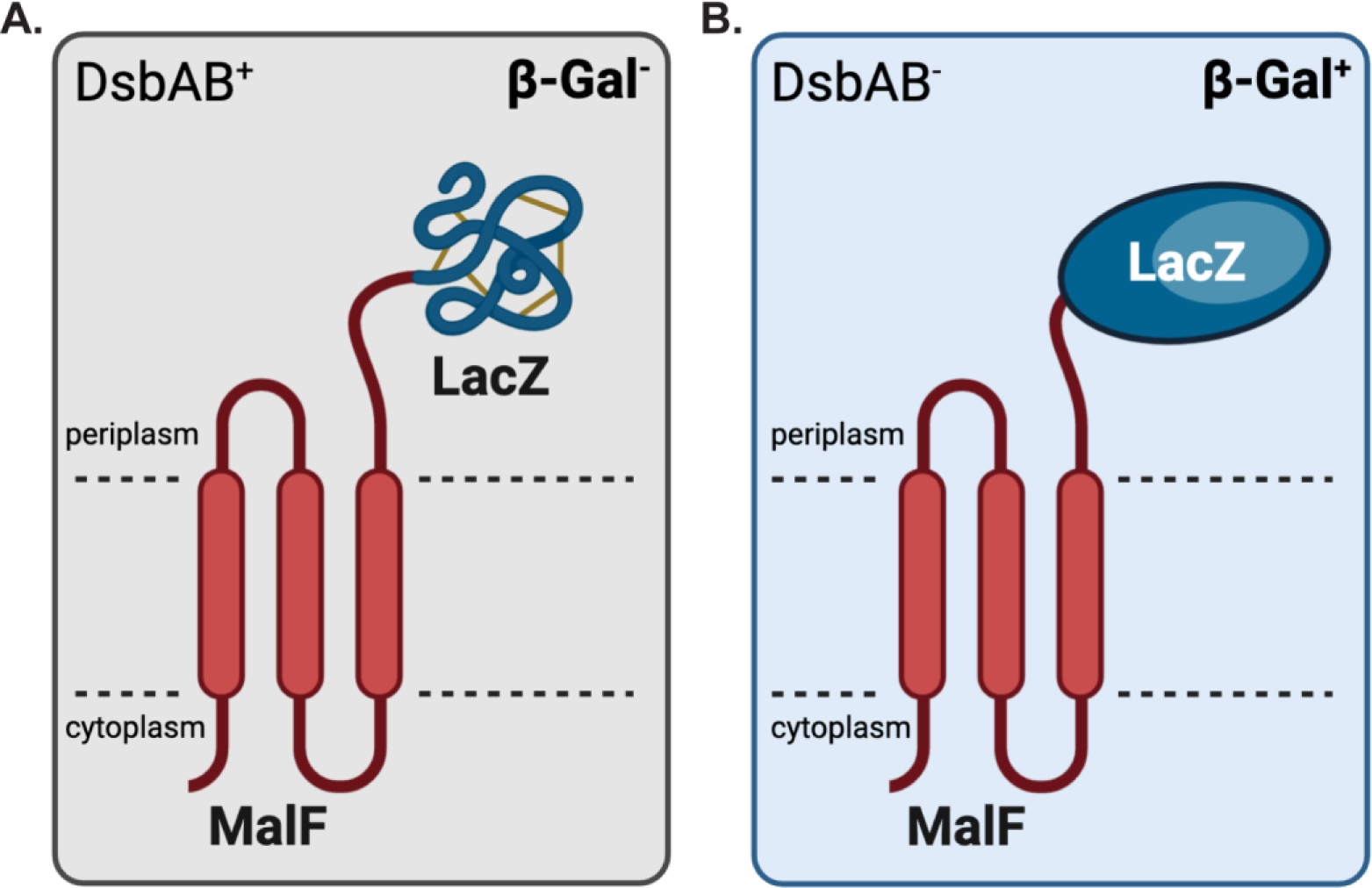
β-Gal^dbs^ (MalF-LacZ102) monitors disulfide bond formation in the bacterial periplasm. A) In *E. coli*, DsbA and DsbB participate in introducing aberrant disulfide bonds (represented as yellow lines) into periplasmic β-Galactosidase hence misfolding and inactivating it. B) The removal or inhibition of either *dsbA* or *dsbB* disrupts the formation of disulfide bonds allowing β-Galactosidase to be folded.

Disulfide bond formation plays a role in protein folding for both eukaryotes and prokaryotes. In bacteria, disulfide bonds contribute to the folding and stability of proteins involved in crucial cellular processes in the cell envelope (6–8). In *E. coli*, disulfide bond formation is catalyzed by DsbA and DsbB enzymes, which work together to introduce disulfide bonds into many proteins. DsbA is a periplasmic protein, a member of the thioredoxin family, that catalyzes the formation of disulfide bonds into substrate proteins through its Cys-X-X-Cys active site (2). DsbB is a membrane protein that regenerates DsbA’s activity by transferring electrons to quinones (9–12). Some pathogenic bacteria, like *Pseudomonas aeruginosa*, harbor more than one copy of the DsbA-DsbB proteins (7, 13). While actinobacteria, cyanobacteria, and δ-proteobacteria, use an alternative enzyme named VKOR (for <u>v</u>itamin <u>K</u> ep<u>o</u>xide <u>r</u>eductase) that performs the same function as DsbB (14). DsbB and VKOR share no protein sequence identity, but they exhibit similar structural features and contain a quinone cofactor to generate a disulfide bond (15, 16).

The viability of mutants lacking disulfide bond-forming machinery varies from organism to organism. *E. coli dsb* mutants are viable aerobically but not anaerobically (17). Overall, disulfide bond formation is required for virulence but not for *in vitro* growth of gram-negative bacteria (6, 7), whereas it is essential in actinobacteria (18–20). Disulfide bond-forming enzymes represent a compelling new drug target because their inhibition could simultaneously affect several proteins localized in the cell envelope, including virulence factors, proteins involved in outer membrane biogenesis, cell division, and antibiotic resistance (6, 7, 21–23). Thus, we have previously developed a cell-based and target-based assay to find molecules that inhibit the membrane proteins, DsbB and VKOR, of pathogenic bacteria (24, 25). This assay uses *E. coli* Δ*dsbB* mutant expressing the β-Gal^dbs^ in which *dsbB* function is complemented with a plasmid carrying either *dsbB* or *vkor* genes from pathogens (24). We then perform parallel screens of compounds to look for inhibitors of each of the enzymes (24–26). The screens are performed in parallel to provide reciprocal controls that eliminate inhibitors that influence β-Gal activity by acting directly on *E. coli* DsbA or affecting membrane protein assembly since those molecules would likely appear as inhibitors of both DsbB and VKOR-expressing strains. We have identified several classes of DsbB and VKOR inhibitors using this method (24, 25, 27) as we have sought to expand the small molecule search by using cell-based and target-based approaches with gram-negative organisms for which antimicrobial resistance is posing a threat.

Here we report the construction of a series of biosensor plasmids that allow fine-tuning the expression of β-Gal^dbs^ and are mobilizable into other gram-negative bacteria. A plasmid carrying β-Gal^dbs^ introduced into *P. aeruginosa* UCBPP-PA14 behaves similarly as in *E. coli*. The strains expressing β-Gal^dbs^ respond to molecular inhibition of the two DsbB proteins and provide a basis to look for novel inhibitors of both enzymes using *P. aeruginosa* cells. These vectors represent tools to further study oxidative protein folding in other gram-negative pathogens and to identify inhibitors of DsbAB proteins against these organisms.

## Results

### Use of tightly regulated promoters to control β-Gal^dbs^ expression

Previous attempts to move the β-Gal^dbs^ into other organisms including *P. aeruginosa* were unsuccessful because moving the plasmid resulted in toxicity which prevented growth of transconjugants. Dwyer *et al*., have shown that periplasmic LacZ is toxic to *E. coli*, likely due to the presence of 16 cysteines which cause aberrant disulfide bonds introduced by DsbA rendering misfolded β-Gal (5). The original construct (Figure 2A, pNG102) includes the maltose-binding protein, MalE, and β-Gal^dbs^ under the maltose promoter (P*_mal_*) (1) which was then inserted into the chromosome at the lambda *att* site (28). We reasoned that the difficulty in introducing the β-Gal^dbs^ construct was perhaps a combination of the basal expression levels and/or the co-expression of MalE upstream of β-Gal^dbs^. We thus cloned only β-Gal^dbs^ into four low-copy number plasmids (p15A ori) with promoters and regulators that have been engineered to have low background, low cross reactivity and high dynamic range (29). The promoters are inducible with cuminic acid (Cuma), vanillic acid (Van), 2,4-diacetylphophloroglucinol (DAPG), anhydrotetracycline (aTc).

**Figure 2.**
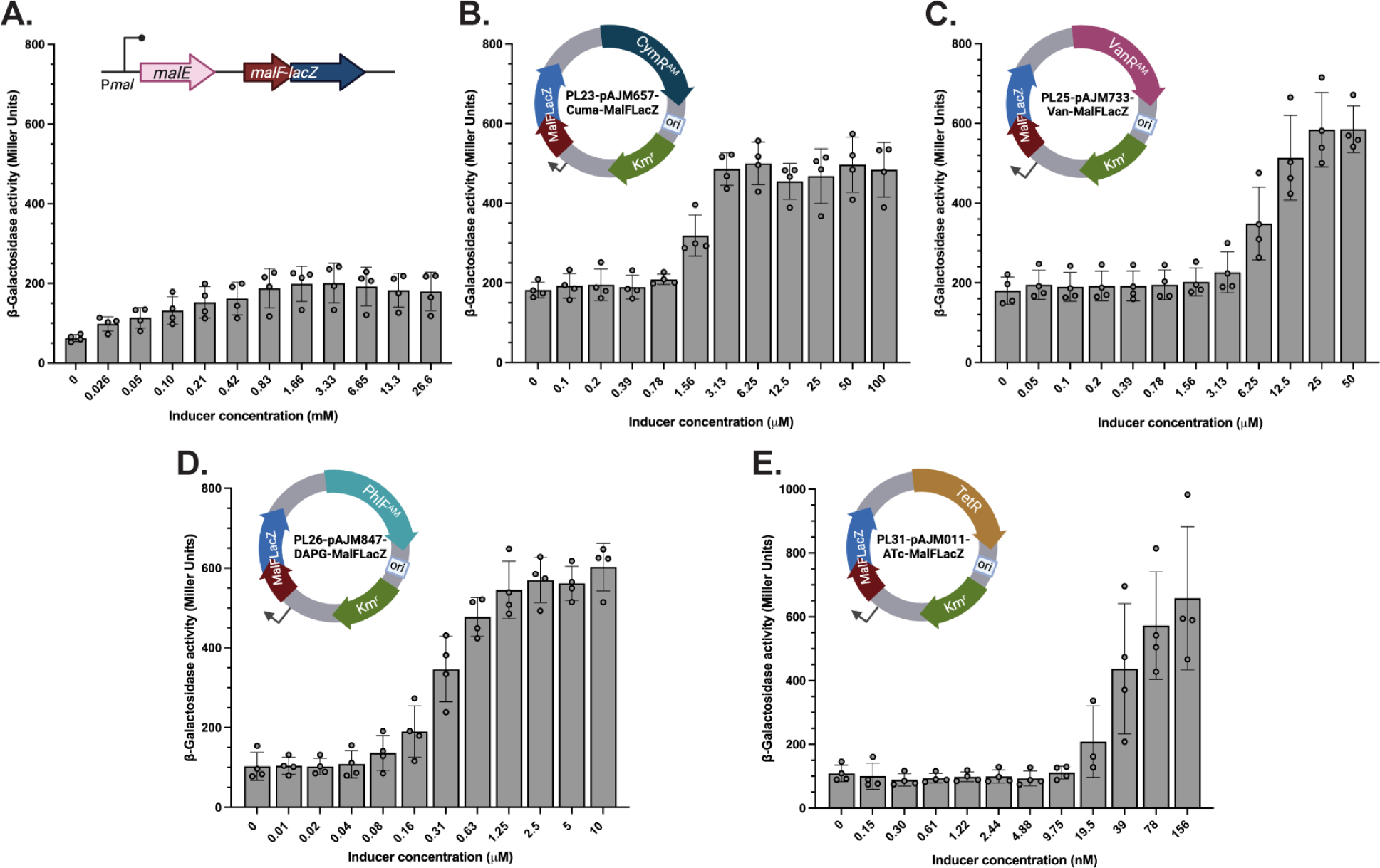
Fine-tuning levels of β-Gal^dbs^ sensor in *E. coli*. A) β-Gal activity of the λ::P_mal_-MalE-β-Gal^dbs^ expressed in *E. coli* Δ*dsbB* strain grown in M63 0.2% glucose supplemented with 50 μg/mL of essential amino acids (except Cys) grown at 30 °C for 10 h and induced with varying concentrations of maltose. An aliquot of cells was then used to quantify β-Gal by following the hydrolysis of ONPG at 28 °C using whole cells. B) β-Gal activity of the P_CymR_-β-Gal^dbs^ expressed in *E. coli* Δ*dsbB* strain induced with Cuma. C) β-Gal activity of the P_VanR_-β-Gal^dbs^ expressed in *E. coli* Δ*dsbB* strain induced with Van. D) β-Gal activity of the P_PhlF_-β-Gal^dbs^ expressed in *E. coli* Δ*dsbB* strain induced with DAPG. E) β-Gal activity of the P_Tet_-β-Gal^dbs^ expressed in *E. coli* Δ*dsbB* strain induced with aTc. Data represents average±SD of four independent experiments. Plasmid maps were created with BioRender.com.

We transformed the plasmids (PL23, PL25, PL26, PL31) into *E. coli* Δ*dsbB* mutant in which β-Gal^dbs^ would be active and quantified the β-Gal activity in M63 minimal medium supplemented with 0.2% glucose, 50 µg/mL of all essential amino acids except cysteine to prevent thiol rearrangement, and serial dilutions of inducers. We used whole cells to measure the velocity of o-nitrophenyl-β-galactoside hydrolysis without lysing cells (no chloroform-SDS step) since ONPG while unable to enter the cytoplasm, is permeable to the outer membrane, and β-Gal^dbs^ is in the periplasm. All four plasmids (Figure 2B-2E) impart higher β-Gal activity compared to P*_mal_*-β-Gal^dbs^ induced with maltose (Figure 2A). To avoid overexpression and toxicity from P*_mal_*-β-Gal^dbs^, we mildly induced adding maltose in media containing glucose to weaken the expression of β-Gal (Figure 2A). The four new vectors allow titration of β-Gal^dbs^ expression at desired levels to sensitize or strengthen the assay (Figure 2). In addition, all the concentrations used to induce the β-Gal^dbs^ resulted in similar growth compared to the growth obtained in strains harboring the P*_mal_*-β-Gal^dbs^ (Supplementary Figure 1). Expression of the aTc-inducible β-Gal^dbs^ vector (PL31), the biosensor plasmid with the highest expression, we see a decrease in growth of the Δ*dsbB* mutant only at very high induction with 10 µM aTc (Supplementary Figure 2).

### Expression of β-Gal^dbs^ in *P. aeruginosa* UCBPP-PA14

*P. aeruginosa* is one of the six pathogens for which new antibacterial agents are most desperately needed (30, 31) and is often associated with extremely difficult-to-treat infections in immunosuppressed patients for whom current antibiotic treatments fail to work (32–35). Recalcitrant infections include acute pneumonia, ulcerative keratitis, bacteremia, urinary tract, intra-abdominal, chronic airway, and wound infections (32, 36).

Some *P. aeruginosa* virulence factors known to contain disulfide bonds include the pilin protein PilA (37, 38), the elastolytic metalloprotease elastase (LasB), exotoxin A (39–41), the chitin-binding protein (CbpD), the immunomodulating metalloprotease IMPa and proteases PaAP and PrpL as well as type-III secretion components required for the delivery of the effectors ExoT and ExoU (37). Disulfide bond-forming enzymes in *P. aeruginosa* include two *dsbA* and two *dsbB* homologs. Disulfide bond formation is mainly driven by DsbA1 which is oxidized by both DsbB1 and DsbB2 proteins (13, 25, 37). The deletion of *dsbA1* and the double deletion of *dsbB1dsbB2* cause a decrease in virulence in pneumonia and keratitis mice models (25, 42). Thus, inhibitors of the disulfide bond-forming enzymes would affect the folding of several virulence factors.

To use the β-Gal^dbs^ biosensor system in other gram-negative bacteria, we subcloned the four regulators and promoters together with β-Gal^dbs^ into a pJN105, a broad host range (BHR) plasmid. pJN105 plasmid harbors a gentamicin resistance cassette, a BHR origin of replication (pBBR) (43), and a mob site to allow conjugal delivery from *E. coli* strains harboring RK2 transfer machinery (44). We then moved the Cuma-inducible plasmid (PL60) into *P. aeruginosa* UCBPP-PA14 wildtype as well as the single and double *dsbB1* and *dsbB2* mutant strains. Both *P. aeruginosa* DsbB proteins maintain oxidized DsbA1, hence the single *dsbB* deletions are rescued by the remaining paralog and reactivate DsbA (13, 25). Indeed, high levels of β-Gal can be observed when we induce the expression of β-Gal^dbs^ with 5 or 50 μM Cuma in the Δ*dsbB1*Δ*dsbB2* mutant grown in M63 defined medium (0.2% glucose) supplemented with 50 μg/mL of all essential amino acids except cysteine (Figure 3B). However, little to no activity is seen in the wildtype or single Δ*dsbB1* or Δ*dsbB2* mutants (Figure 3B). Furthermore, induction of β-Gal^dbs^ displays a good range over increasing concentrations of Cuma and there is also lower background activity in the absence of the inducer compared to what is observed in *E. coli* (Figure 3C). Thus, the concentration of the inducer can be adjusted to the desired sensitivity of the assay when screening for inhibitors of the Dsb proteins (see below).

**Figure 3.**
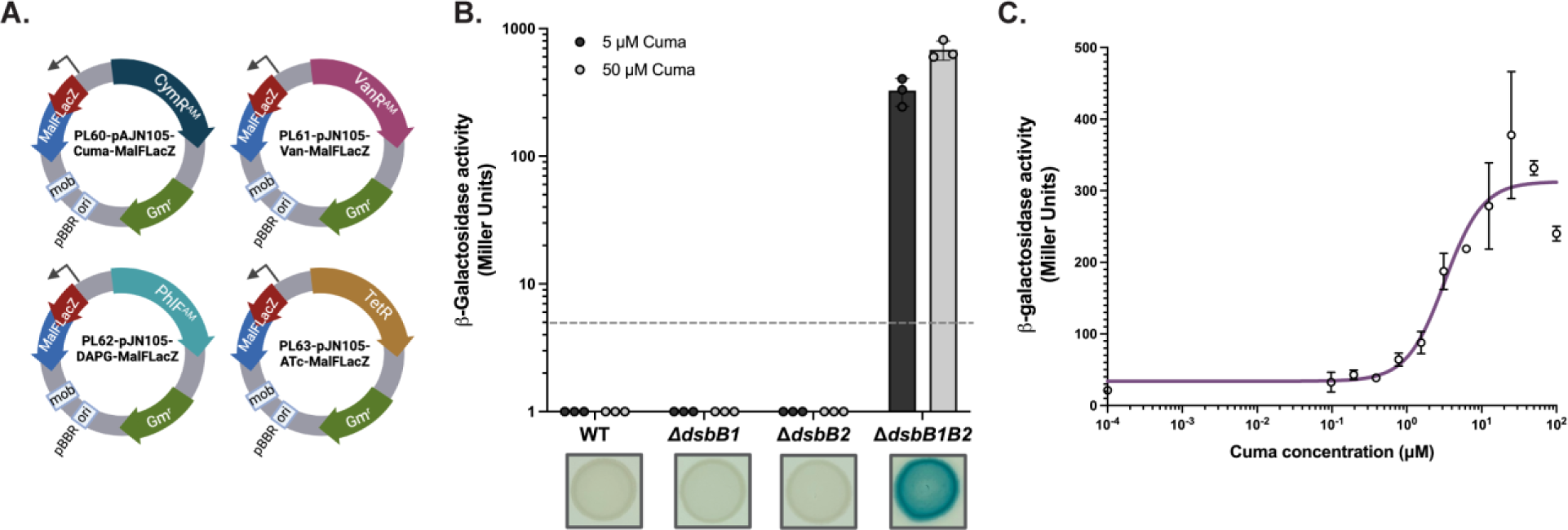
β-Gal^dbs^ sensors for gram-negative bacteria. A) Mobilizable vectors carrying β-Gal^dbs^ under regulatable promoters. B) Low levels of β-Gal^dbs^ (limit of detection: 5 MU) in wildtype, Δ*dsbB1* and Δ*dsbB2* where DsbA is still functional, while the double Δ*dsbB1*Δ*dsbB2* mutant allows activation of β-Gal due to lack of DsbA oxidation. PL60 was conjugated into *P. aeruginosa* UCBPP-PA14 wildtype and *dsbB* mutants. β-Gal^dbs^ was measured by ONPG hydrolysis in live cells that were grown for 18 h at 37 °C in M63 0.2% glucose supplemented with 50 μg/mL of essential amino acids (except Cys), antibiotics and either 5 or 50 μM of Cuma to induce β-Gal^dbs^. Data represents average±SD of three independent experiments. 10 μL aliquots of overnight cultures were spotted onto M63 0.2% glucose supplemented with 50 μM Cuma and 120 μg/mL X-Gal. Plates were incubated at 30 °C for 2 days. Images represent the result of at least three independent experiments. C) The β-Gal activity is dose-dependent on the amount of Cuma inducer in the double Δ*dsbB1*Δ*dsbB2* mutant. β-Gal^dbs^ was measured as in B) but with a serial dilution of Cuma to induce β-Gal^dbs^. Data represents average±SD of three independent experiments and non-linear regression was used to model the response. Plasmid maps were created with BioRender.com.

### Inhibition of DsbB1 and DsbB2 using *P. aeruginosa* β-Gal^dbs^ sensor

As a proof of concept for a small molecule screen, we tested the Δ*dsbB1* and Δ*dsbB2* mutants against an inhibitor of the two proteins that we previously discovered, compound 12, a dichloro-pyridazinone (24). We used both agar (Figure 4A) and liquid (Figure 4B) assays to determine the inhibition of DsbB1 and DsbB2 proteins independently. The adaptation of a 384-well plate agar assay using X-Gal, for the Δ*dsbB1* mutant gave a dark blue color at high concentrations of compound 12 when β-Gal^dbs^ was induced at 5 μM Cuma (Figure 4A, left). A higher concentration of β-Gal^dbs^ inducer, 25 μM Cuma, was needed for the Δ*dsbB2* mutant to show observable activity, with even high concentrations of compound 12 only causing light blue pigmentation (Figure 4A, right). We were also able to adapt a 96-well plate liquid assay from the *E. coli* assay by growing *P. aeruginosa* in M63 defined medium supplemented with 50 μg/mL of all essential amino acids except cysteine. As with the *E. coli* assay, the ONPG hydrolysis was measured without lysing *P. aeruginosa* cells. The quantification of β-Gal activity (Figure 4B) reflected what was observed in the agar plates, the Δ*dsbB1* mutant displays increased β-Gal activity at higher concentrations of compound 12 than the Δ*dsbB2* mutant. This is consistent with our previous findings using indirect assays for inhibition of DsbB proteins by measuring elastase activity in the supernatant of *P. aeruginosa* grown with the drug (25). While compound 12 behaves as a strong inhibitor in an *E. coli* strain expressing *P. aeruginosa* DsbB1, when using *P. aeruginosa* the drug behaves as a stronger inhibitor of DsbB2 (Δ*dsbB1* strain) than DsbB1 (Δ*dsbB2* strain) and the drug does not reach full inhibition of elastase when the DsbB1 protein is present (Supplementary Figure 3). This divergence between hosts was one of the motivations for the development of a target-based whole cell-based method using the native host, in this case, *P. aeruginosa* itself.

**Figure 4.**
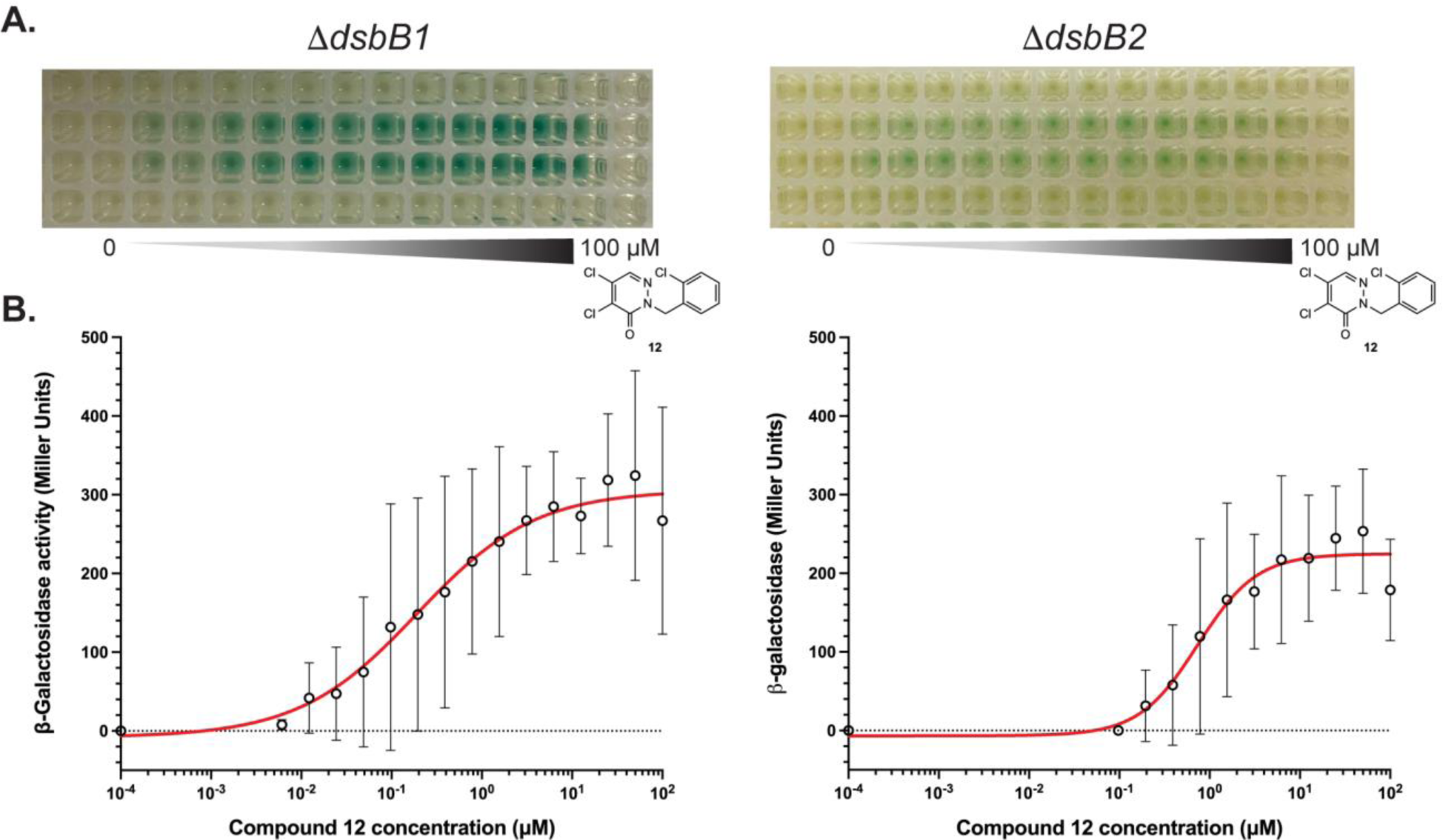
β-Gal^dbs^ expression allows detection of DsbB inhibition in *P. aeruginosa*. A) Inhibition of DsbB2 (Δ*dsbB1*, left) and DsbB1 (Δ*dsbB2,* right) with compound 12 can be observed by the generation of blue color when expressing β-Gal^dbs^ in the mutants. 384-well plates were prepared with 50 μL of M63 0.2% glucose supplemented with X-Gal and either 5 (Δ*dsbB1*) or 25 μM (Δ*dsbB2*) Cuma to induce β-Gal^dbs^. A two-fold serial dilution of 6 mM compound 12 was made in DMSO and 1 μL of the diluted drug was pipetted into the agar surface. 10 μL of bacteria were then added on top and incubated for 2 days at 30 °C. Images represent the result of at least two independent experiments. B) Quantification of β-Gal in Δ*dsbB1* and Δ*dsbB2* mutants grown in the presence of compound 12. β-Gal^dbs^ was measured by ONPG hydrolysis in live *P. aeruginosa dsbB* cells that were previously grown for 18 h at 37 °C in M63 0.2% glucose supplemented with 50 μg/mL of essential amino acids (except Cys), 5 μM (Δ*dsbB1*) or 25 μM (Δ*dsbB2*) Cuma, and serially-diluted compound 12. Data represents average±SD of three independent experiments and non-linear regression was used to model the response.

## Discussion

The MalF-LacZ102 hybrid protein (β-Gal^dbs^) has been an instrumental tool in studying disulfide bond formation in *E. coli* for over three decades. The use of LacZ allows for simple genetic selections and screens (3). Basal expression of the hybrid protein in M63 glucose minimal medium without maltose allowed the development of a heterologous platform to search for inhibitors of DsbB and VKOR proteins via high throughput screening using *E. coli* (24, 25). However, the misfolding of β-Gal^dbs^ in the periplasm results in toxicity and cell death due to aberrant disulfide bonds formed by DsbA when the fusion was highly induced from *P_mal_-*β-Gal^dbs^ in the presence of maltose and glycerol in the medium (5). Indeed, our previous attempts to use the β-Gal^dbs^ in other gram-negative organisms failed likely due to this toxicity problem. In this work, we showed that it is possible to express the β-Gal^dbs^ in a heterologous host such as, *P. aeruginosa*, when using tightly regulated promoters. The expression of β-Gal^dbs^ under the Cuma-inducible promoter was tolerated in *P. aeruginosa* cells and no growth defect was observed at the maximum inducer concentration tested. Deletion of *dsbA* or both *dsbB* proteins in *P. aeruginosa* cells expressing the β-Gal^dbs^ allows the folding of periplasmic β-Gal and hence produces blue colonies in the presence of X-Gal, similar to what is observed for the single deletions of *dsbA* or *dsbB* in *E. coli*. We were also able to adapt our *E. coli* high (384-well plate) and medium-throughput (96-well plate) methods expressing the β-Gal^dbs^ sensor in *P. aeruginosa.* The Cuma-inducible promoter gave a good dynamic range to look for inhibitors of Dsb proteins in *P. aeruginosa*. The amount of inducer can be optimized to a desired level to sensitize the assay. For instance, with more β-Gal^dbs^ expressed to a level where there is already a pale blue color (when DsbAB cannot misfold all β-Gal^dbs^ produced) one would detect weaker inhibitors, while less expression of the fusion would find stronger inhibitors (24).

Considering that disulfide bonds are required for the stability of virulence factors, toxins, antibiotic resistance, and cell envelope biogenesis proteins, lack of disulfide bond formation in many gram-negative pathogens results in virulence attenuation as well as antibiotic and phage susceptibility (6, 7, 21, 23, 45). However, the pathway is not essential for aerobic laboratory growth (17), thus facilitating the development of methods to search for molecules that identify molecules on-target in pathogenic bacteria, also known as target-based and whole cell-based approaches (46). The biosensor plasmids generated in this work could not only help to identify such Dsb inhibitors but also to study the variations of disulfide bond formation found in other bacteria (7, 14). Some gram-negative organisms encode two or more sets of DsbAB proteins that seem to play different roles but can substitute for each other in oxidizing particular substrates. Examples of organisms with multiple Dsb proteins include *P. aeruginosa* (13), *Salmonella enterica* serovar Typhimurium (47), *Neisseria meningitidis* (48, 49), and *Campylobacter jejuni* (50). In addition, some proteobacterial DsbA proteins such as *Legionella pneumophila* DsbA can perform both reduction and oxidation of disulfide bonds (51). The redundancy of Dsb proteins could perhaps provide a specialized activity for folding certain substrates that require priority under some environmental conditions. Using the single *dsb* mutants expressing the β-Gal^dbs^ one could look for an array of growth conditions that inactivate the Dsb protein and render Lac+ cells. Similarly, one could also look for intragenic *L. pneumophila dsbA* mutations to either loose or gain a more oxidizing nature with the use of the β-Gal^dbs^ by looking for pale blue or white colonies, respectively. In addition, the β-Gal^dbs^ can be used to study bacterial interactions between organisms that share the same niche, such as the gut to determine whether other bacteria could affect the oxidative protein folding of their neighbors through secretion of quinones (52) or other redox molecules.

The use of translational LacZ fusions has advanced the study of membrane topology, protein targeting and secretion as well as oxidative protein folding in *E. coli* since the 70’s (3). The use of the BHR plasmid-borne β-Gal^dbs^ disulfide bond biosensor we describe here should enable this analysis in a greater diversity of bacteria.

## Acknowledgments

To my dear friend and mentor Jon Beckwith, for sharing your genius, your vision, your wisdom, and your generosity. We thank Laura McPartland for her help in constructing pLEM2. Drawings in Figure 1, Figure 2, and Figure 3A were created using Biorender.com with publication licenses KF265H9JZ7 and BX265SG74U. We also thank Clay Fuqua for helpful suggestions and comments on this manuscript. This work was supported by Indiana University Bloomington and by Cystic Fibrosis Foundation Pilot and Feasibility Award 004846I222 (to C.L.).

## Methods

### Strains and growth conditions

The strains and plasmids used in this study are listed in Table 1. To construct PL23, 25, 26 and PL31, plasmid backbones without eYFP were amplified by PCR with PR41 and PR42 using Q5 Polymerase (New England Biolabs) and pAJM657, pAJM773, pAJM847 and pAJM011 as templates. The β-Gal^dbs^ insert was amplified with PR43 and PR44 using HK325 genomic DNA as a template. Insert and vectors were independently assembled using BsaI-HFv2 Golden Gate Assembly (NEBridge, New England Biolabs). Ligated products were transformed into HK320 electrocompetent cells and plated on NZ Kanamycin, inducer (Cuma, Van, DAPG, or aTc) and X-Gal. One blue colony was used to purify the plasmid and sequence the insert using primers PR51 and PR52. To construct PL60-PL63, high-fidelity assembly (NEBuilder, New England Biolabs) was used using three PCR products. First, PR96 and PR97 were used to amplify the backbone with β-Gal^dbs^ without *araC* and arabinose promoter using pLEM2 vector. Second, the four regulators were amplified with PR98 and PR99 while the four promoters were amplified with PR100 and PR101 using pAJM657, pAJM773, pAJM847 and pAJM011 as templates. The three inserts were assembled independently using NEBuilder and transformed into DH10β competent cells (New England Biolabs). Plasmids were sequenced with PR102 and PR46 to confirm the regulators and promoters. PL60 was then transformed into *E. coli* S17λpir electrocompetent cells and used to conjugate the vector into *P. aeruginosa* strains. Nalidixic acid was used to counter select the *E. coli* donor and gentamicin to select for the plasmids.

**Table 1.**
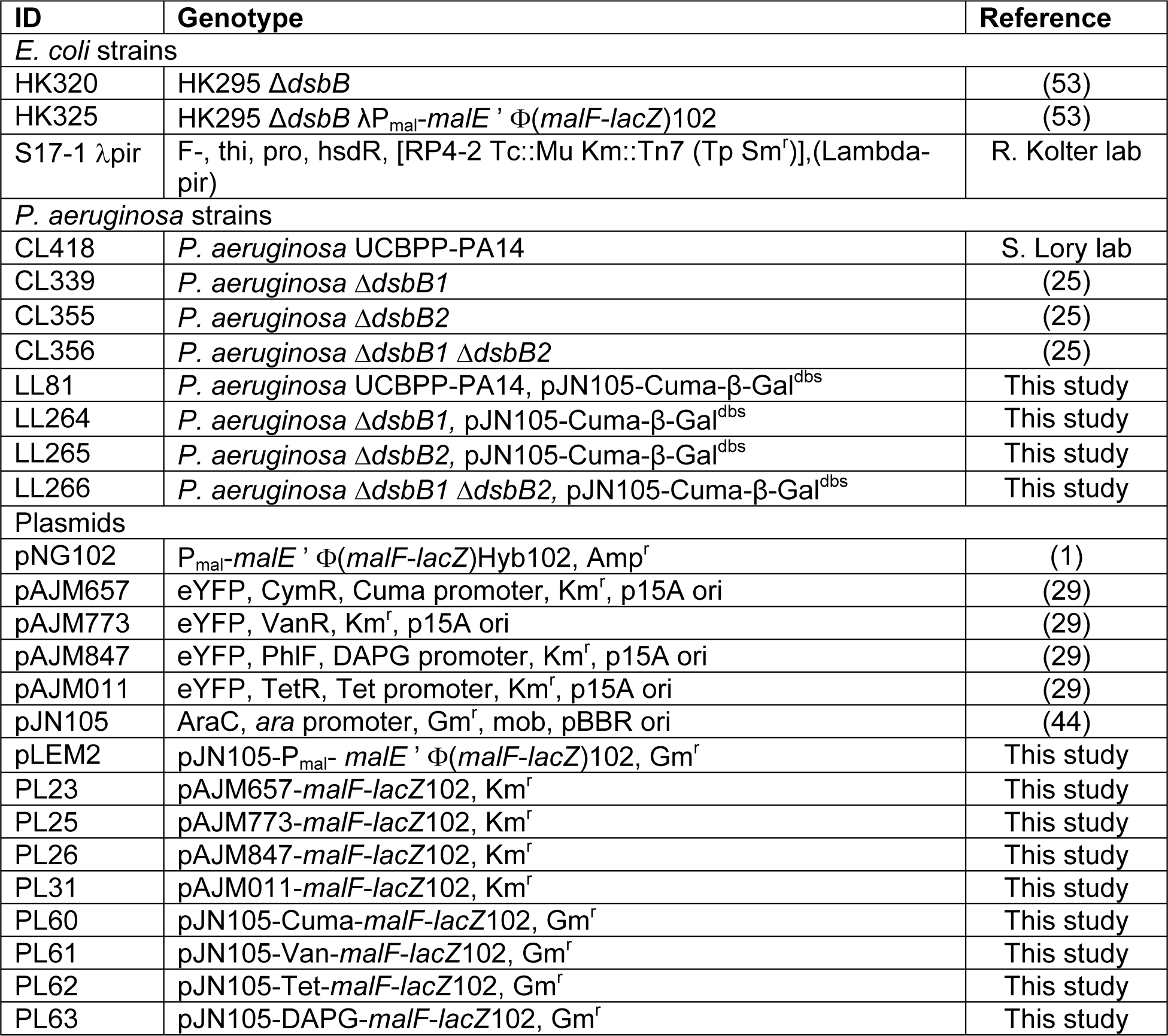
List of strains.

**Table 2.**
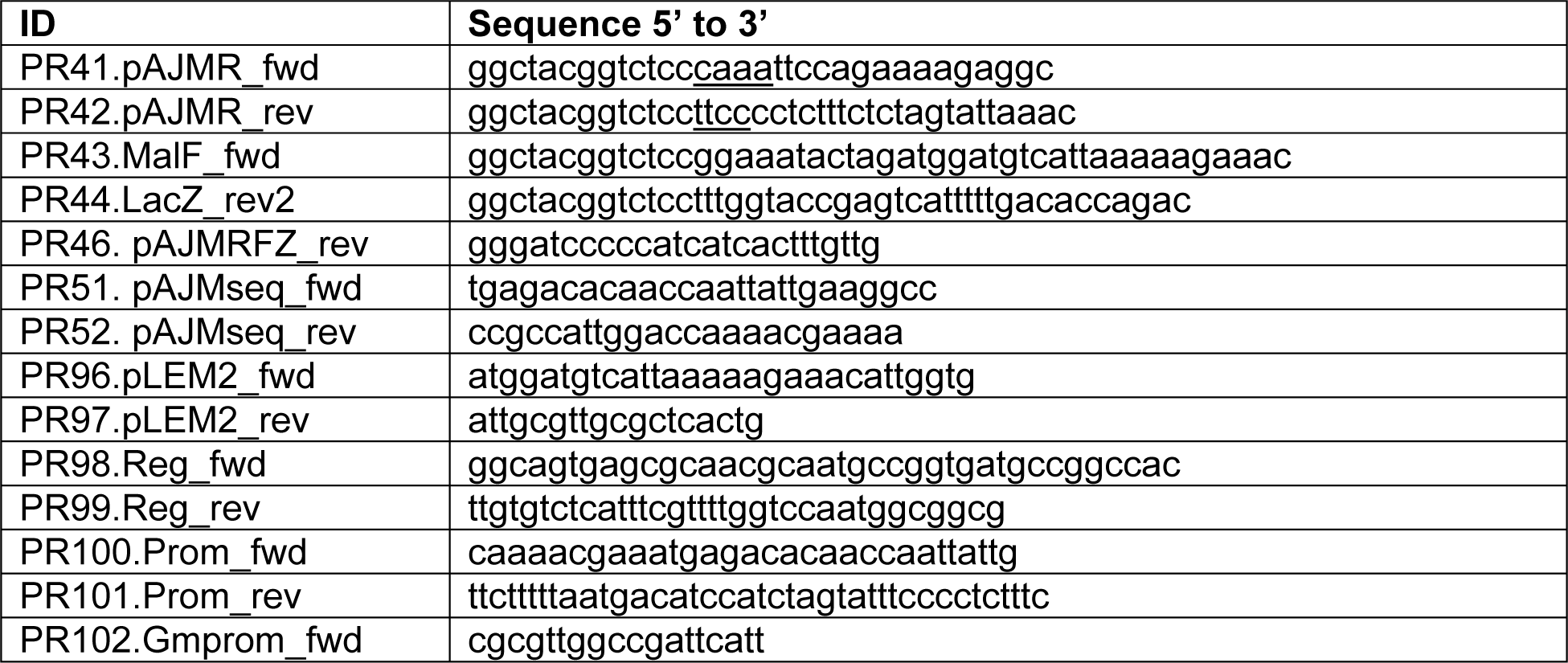
Primer list.

All *E. coli* strains were grown in NZ or M63 0.2% glucose and supplemented with 50 μg/mL of 19 essential amino acids (Ser, Val, Ile, Leu, Trp, Tyr, Met, Asp, Glu, Ala, Arg, Lys, Asn, Gln, Phe, Gly, Thr, His and Pro) at 30-37°C when indicated. All *P. aeruginosa* strains were grown in LB or M63 0.2% glucose supplemented with 50 μg/mL of 19 essential amino acids either broth or agar media at 30-37°C when indicated. The antibiotic concentrations used were nalidixic acid 10 μg/mL, kanamycin 40 μg/mL, gentamicin 3 μg/mL for *E. coli* and 10 μg/mL for *P. aeruginosa*.

Kanamycin was purchased from Sigma (USA). Nalidixic acid and gentamicin were purchased from GoldBio (USA). Inducers were purchased from Sigma (Cuminic acid, Vanillic acid, and anhydrotetracycline hydrochloride) and ChemCruz (2,4-Diacetylphlorogucinol). Molecules dissolved in DMSO to make 100 mM (Cuma, Van, DAPG) and 1 mM (aTc) stocks, respectively. Compound 12 was purchased from Enamine (EN300-173996 purity 95%, Ukraine) and dissolved in DMSO to 10 mg/mL (34.72 mM).

### β-Galactosidase assays

β-Galactosidase assays were done by determining the velocity of hydrolysis of o-nitrophenyl-β-galactoside (ONPG, Sigma) in microtiter plates using a protocol previously described with slight modifications (24, 27, 54). Briefly, cultures were grown in M63 0.2% glucose medium with proper antibiotics at 37°C overnight. Cells were then inoculated to an OD600 of 0.01 into fresh M63 medium containing 0.2% glucose, 50 μg/mL of 19 essential amino acids (except cysteine) and proper antibiotics. 200 μL of diluted bacteria were transferred to a 96-well plate (Corning) and a two-fold serial dilution of the inducers was done from columns 12 to 2. The plate was then sealed with a breathable film. The growth plate was incubated for 10 h at 30°C and fast-orbital shaking inside a Synergy H1 plate reader (BioTek). After growth, absorbance at 600nm was read in a Synergy H1 plate reader (BioTek). Then, 100 μL of bacteria from the growth plate were transferred to the assay plate. Note that no cell lysis step is required because β-Gal^dbs^ is in the periplasm. The reaction was started by adding 100 μL of the ONPG buffer to the cells (a mixture of 8 mL of Z-buffer with 4 mL of 4 mg/mL ONPG dissolved in Z-buffer). The addition of β-mercaptoethanol was also omitted to avoid interference with the disulfide bonds formed in β-Gal^dbs^. The absorbance at λ420nm was measured every minute for 1-2 h to follow the kinetics of ONPG hydrolysis at 28 °C in a Synergy H1 plate reader (BioTek). The velocity of the reaction was calculated by performing linear regression using GraphPad Prism software. The slopes were then used together with OD600 and the following constants 1.81 (CF1), 2.45 (CF2), and 3.05 (CF3) to calculate Miller Units as reported before (54). *P. aeruginosa* β-Galactosidase assays were done in microtiter plates like the *E. coli* protocol with few modifications in the growth conditions. Cultures were grown in M63 0.2% glucose medium supplemented with 100 μL of NZ broth and proper antibiotics at 37°C overnight. Cells were then inoculated to an OD600 of 0.01 into fresh M63 medium containing 0.2% glucose, 50 μg/mL of 19 essential amino acids (except cysteine), proper antibiotics, and either a fixed concentration or a serial dilution of Cuma when indicated. 200 μL of diluted bacteria were transferred to a 96-well plate (Corning). 100 μM compound 12 was added to column 12 and two-fold serially diluted to column 2. Plate was sealed with a breathable film and incubated for 18 h at 37 °C and 700 rpm in an Incu-mixer MP (Benchmark). After growth, the following steps were done like the *E. coli* assay indicated above. Non-linear regression using GraphPad Prism Software (Boston, USA) was used to model the data by using the equation [Agonist] vs. response - variable slope (four parameters).

### Agar assays

Drug testing was performed as previously described with slight modifications (24, 25). A liquid dispenser (BioTek) fitted with a small-bore tubing cartridge was used to dispense 50 μL aliquots of hot agar medium [M63 medium containing 0.2% glucose and 0.9% agar, supplemented with gentamicin (10 μg/mL), Cuma (5 or 25 μM), and X-Gal (120 μg/mL)] to 384-well tissue culture-treated plates (BD Falcon). In order to prevent agar solidification in the Wellmate tubing (at too-low temperatures) or inactivation of the antibiotics and X-Gal (at too-high temperatures), the medium was maintained in a 60 °C oven. In addition, the Wellmate tubing was pre-warmed by washing with sterile hot water immediately prior to loading the agar medium. After the agar solidified, the plates were stored in a humidified sealed container at 4 °C for no more than 2 days. Compound 12 was added by pipetting 1 μL of serial dilutions into the agar’s surface (final concentration of DMSO, 1.7%) and 10 μL of diluted *P. aeruginosa* cells (OD600 of 0.01) were dispensed with a liquid dispenser (BioTek). 384-well plates were sealed with a breathable film and then incubated in humidity boxes at 30°C for two days and then stored at 4°C for two days to determine the minimal concentration to produce a blue color. Photographs were taken using a white LED transilluminator.

## References

1. Froshauer S, Green GN, McGovern DBK, Beckwith J. 1988. Genetic analysis of the membrane insertion and topology of MalF, a cytoplasmic membrane protein of Escherichia coli. J Mol Biol 200:501–511.

2. Bardwell JCA, McGovern K, Beckwith J. 1991. Identification of a protein required for disulfide bond formation in vivo. Cell 67:581–589.

3. Beckwith J. 2013. Fifty years fused to lac. Annu Rev Microbiol 67:1–19.

4. Manoil C, Mekalanos JJ, Beckwith2 J. MINIREVIEW Alkaline Phosphatase Fusions: Sensors of Subcellular Location.

5. Dwyer RS, Malinverni JC, Boyd D, Beckwith J, Silhavy TJ. 2014. Folding LacZ in the periplasm of Escherichia coli. J Bacteriol 196:3343–3350.

6. Heras B, Shouldice SR, Totsika M, Scanlon MJ, Schembri MA, Martin JL. 2009. DSB proteins and bacterial pathogenicity. Nat Rev Microbiol 7:215–25.

7. Landeta C, Boyd D, Beckwith J. 2018. Disulfide bond formation in prokaryotes. Nat Microbiol 3:270–280.

8. Collet JF, Cho SH, Iorga BI, Goemans C V. 2020. How the assembly and protection of the bacterial cell envelope depend on cysteine residues. Journal of Biological Chemistry 295:11984–11994.

9. Bardwell JC, Lee JO, Jander G, Martin N, Belin D, Beckwith J. 1993. A pathway for disulfide bond formation in vivo. Proc Natl Acad Sci U S A 90:1038–42.

10. Missiakas D, Georgopoulos C, Raina S. 1993. Identification and characterization of the Escherichia coli gene dsbB, whose product is involved in the formation of disulfide bonds in vivo. Proc Natl Acad Sci U S A 90:7084–8.

11. Bader M, Muse W, Ballou DP, Gassner C, Bardwell JCA. 1999. Oxidative protein folding is driven by the electron transport system. Cell 98:217–227.

12. Kadokura H, Beckwith J. 2002. Four cysteines of the membrane protein DsbB act in concert to oxidize its substrate DsbA. EMBO Journal 21:2354–2363.

13. Arts IS, Ball G, Leverrier P, Garvis S, Nicolaes V, Vertommen D, Ize B, Dufe VT, Messens J, Voulhoux R, Collet J-F. 2013. Dissecting the Machinery That Introduces Disulfide Bonds in Pseudomonas aeruginosa. mBio 4:1–11.

14. Dutton RJ, Boyd D, Berkmen M, Beckwith J. 2008. Bacterial species exhibit diversity in their mechanisms and capacity for protein disulfide bond formation. Proc Natl Acad Sci U S A 105:11933–11938.

15. Inaba K, Murakami S, Nakagawa A, Iida H, Kinjo M, Ito K, Suzuki M. 2009. Dynamic nature of disulphide bond formation catalysts revealed by crystal structures of DsbB. EMBO J 28:779–791.

16. Li W, Schulman S, Dutton RJ, Boyd D, Beckwith J, Rapoport TA. 2010. Structure of a bacterial homologue of vitamin K epoxide reductase. Nature 463:507–12.

17. Meehan BM, Landeta C, Boyd D, Beckwith J. 2017. The disulfide bond formation pathway is essential for anaerobic growth of Escherichia coli. J Bacteriol 199.

18. Dutton RJ, Wayman A, Wei J-R, Rubin EJ, Beckwith J, Boyd D. 2010. Inhibition of bacterial disulfide bond formation by the anticoagulant warfarin. Proc Natl Acad Sci U S A 107:297–301.

19. Ke N, Landeta C, Wang X, Boyd D, Eser M, Beckwith J. 2018. Identification of the thioredoxin partner of VKOR in mycobacterial disulfide bond formation. J Bacteriol 10.1128/jb.00137-18.

20. Reardon-Robinson ME, Osipiuk J, Jooya N, Chang C, Joachimiak A, Das A, Ton-That H. 2015. A thiol-disulfide oxidoreductase of the Gram-positive pathogen Corynebacterium diphtheriae is essential for viability, pilus assembly, toxin production and virulence. Mol Microbiol 98:1037–1050.

21. Furniss RCD, Kadeřábková N, Barker D, Bernal P, Maslova E, Antwi AAA, McNeil HE, Pugh HL, Dortet L, Blair JMA, Larrouy-Maumus G, McCarthy RR, Gonzalez D, Mavridou DAI. 2022. Breaking antimicrobial resistance by disrupting extracytoplasmic protein folding. Elife 11:1–37.

22. Depuydt Matthieu, Messens Joris, Collet Jean-Francois. 2011. How proteins form disulfide bonds. Antioxid Redox Signal 15:49–66.

23. Nikol Kadeřábková, R. Christopher D. Furniss, Evgenia Maslova, Lara Eisaiankhongi, Patricia Bernal, Alain Filloux, Cristina Landeta, Diego Gonzalez, Ronan R. McCarthy, Despoina A.I. Mavridou. 2023. Antibiotic potentiation and inhibition of cross-resistance in pathogens associated with cystic fibrosis. bioRxiv 10.1101/2023.08.02.551661.

24. Landeta C, Blazyk JL, Hatahet F, Meehan BM, Eser M, Myrick A, Bronstain L, Minami S, Arnold H, Ke N, Rubin EJ, Furie BC, Furie B, Beckwith J, Dutton R, Boyd D. 2015. Compounds targeting disulfide bond forming enzyme DsbB of Gram-negative bacteria. Nat Chem Biol 11:292–298.

25. Landeta C, McPartland L, Tran NQ, Meehan BM, Zhang Y, Tanweer Z, Wakabayashi S, Rock J, Kim T, Balasubramanian D, Audette R, Toosky M, Pinkham J, Rubin EJ, Lory S, Pier G, Boyd D, Beckwith J. 2019. Inhibition of Pseudomonas aeruginosa and Mycobacterium tuberculosis disulfide bond forming enzymes. Mol Microbiol 111:918–937.

26. Landeta C, Meehan B, McPartland L, Ingendahl L, Hatahet F, Ngoc T, Boyd D, Beckwith J. 2017. Inhibition of virulence-promoting disulfide bond formation enzyme DsbB is blocked by mutating residues in two distinct regions. J Biol Chem 292:6529–6541.

27. Landeta C, Meehan BM, McPartland L, Ingendahl L, Hatahet F, Tran NQ, Boyd D, Beckwith J. 2017. Inhibition of virulence-promoting disulfide bond formation enzyme DsbB is blocked by mutating residues in two distinct regions. Journal of Biological Chemistry 292:6529–6541.

28. Tian H, Boyd D, Beckwith J. 2000. A mutant hunt for defects in membrane protein assembly yields mutations affecting the bacterial signal recognition particle and Sec machinery. Proc Natl Acad Sci U S A 97:4730–4735.

29. Meyer AJ, Segall-Shapiro TH, Glassey E, Zhang J, Voigt CA. 2019. Escherichia coli “Marionette” strains with 12 highly optimized small-molecule sensors. Nat Chem Biol 15:196–204.

30. Rice LB. 2010. Progress and Challenges in Implementing the Research on ESKAPE Pathogens. Infect Control Hosp Epidemiol 31:S7–S10.

31. Murray CJ, Ikuta KS, Sharara F, Swetschinski L, Robles Aguilar G, Gray A, Han C, Bisignano C, Rao P, Wool E, Johnson SC, Browne AJ, Chipeta MG, Fell F, Hackett S, Haines-Woodhouse G, Kashef Hamadani BH, Kumaran EAP, McManigal B, Agarwal R, Akech S, Albertson S, Amuasi J, Andrews J, Aravkin A, Ashley E, Bailey F, Baker S, Basnyat B, Bekker A, Bender R, Bethou A, Bielicki J, Boonkasidecha S, Bukosia J, Carvalheiro C, Castañeda-Orjuela C, Chansamouth V, Chaurasia S, Chiurchiù S, Chowdhury F, Cook AJ, Cooper B, Cressey TR, Criollo-Mora E, Cunningham M, Darboe S, Day NPJ, De Luca M, Dokova K, Dramowski A, Dunachie SJ, Eckmanns T, Eibach D, Emami A, Feasey N, Fisher-Pearson N, Forrest K, Garrett D, Gastmeier P, Giref AZ, Greer RC, Gupta V, Haller S, Haselbeck A, Hay SI, Holm M, Hopkins S, Iregbu KC, Jacobs J, Jarovsky D, Javanmardi F, Khorana M, Kissoon N, Kobeissi E, Kostyanev T, Krapp F, Krumkamp R, Kumar A, Kyu HH, Lim C, Limmathurotsakul D, Loftus MJ, Lunn M, Ma J, Mturi N, Munera-Huertas T, Musicha P, Mussi-Pinhata MM, Nakamura T, Nanavati R, Nangia S, Newton P, Ngoun C, Novotney A, Nwakanma D, Obiero CW, Olivas-Martinez A, Olliaro P, Ooko E, Ortiz-Brizuela E, Peleg AY, Perrone C, Plakkal N, Ponce-de-Leon A, Raad M, Ramdin T, Riddell A, Roberts T, Robotham JV, Roca A, Rudd KE, Russell N, Schnall J, Scott JAG, Shivamallappa M, Sifuentes-Osornio J, Steenkeste N, Stewardson AJ, Stoeva T, Tasak N, Thaiprakong A, Thwaites G, Turner C, Turner P, van Doorn HR, Velaphi S, Vongpradith A, Vu H, Walsh T, Waner S, Wangrangsimakul T, Wozniak T, Zheng P, Sartorius B, Lopez AD, Stergachis A, Moore C, Dolecek C, Naghavi M. 2022. Global burden of bacterial antimicrobial resistance in 2019: a systematic analysis. The Lancet 399:629–655.

32. Page MG, Heim J. 2009. Prospects for the next anti-Pseudomonas drug. Curr Opin Pharmacol 9:558–565.

33. Maschmeyer G, Braveny I. 2000. Review of the incidence and prognosis of Pseudomonas aeruginosa infections in cancer patients in the 1990s. European Journal of Clinical Microbiology and Infectious Diseases 19:915–925.

34. Fujitani S, Sun HY, Yu VL, Weingarten JA. 2011. Pneumonia due to pseudomonas aeruginosa: Part I: Epidemiology, clinical diagnosis, and source. Chest 139:909–919.

35. Folkesson A, Jelsbak L, Yang L, Johansen HK, Ciofu O, Hoiby N, Molin S. 2012. Adaptation of Pseudomonas aeruginosa to the cystic fibrosis airway: An evolutionary perspective. Nat Rev Microbiol 10:841–851.

36. Lyczak JB, Cannon CL, Pier GB. 2000. Establishment of Pseudomonas aeruginosa infection: Lessons from a versatile opportunist. Microbes Infect 2:1051–1060.

37. Ha U, Wang Y, Jin S. 2003. DsbA of Pseudomonas aeruginosa is essential for multiple virulence factors. Infect Immun 71:1590–1595.

38. Harvey H, Habash M, Aidoo F, Burrows LL. 2009. Single-residue changes in the C-terminal disulfide-bonded loop of the Pseudomonas aeruginosa type IV pilin influence pilus assembly and twitching motility. J Bacteriol 191:6513–6524.

39. Madshus IH, Collier RJ. 1989. Effects of eliminating a disulfide bridge within domain II of Pseudomonas aeruginosa exotoxin A. Infect Immun 57:1873–1878.

40. Braun P, Ockhuijsen C, Eppens E, Koster M, Bitter W, Tommassen J. 2001. Maturation of Pseudomonas aeruginosa elastase: Formation of the disulfide bonds. Journal of Biological Chemistry 276:26030–26035.

41. Urban A, Leipelt M, Eggert T, Jaeger KE. 2001. DsbA and DsbC affect extracellular enzyme formation in Pseudomonas aeruginosa. J Bacteriol 183:587–596.

42. Kim SH, Park SY, Heo YJ, Cho YH. 2008. Drosophila melanogaster-based screening for multihost virulence factors of Pseudomonas aeruginosa PA14 and identification of a virulence-attenuating factor, HudA. Infect Immun 76:4152–4162.

43. Antoine R, Locht C. 1992. Isolation and molecular characterization of a novel broad-host-range plasmid from Bordetella bronchiseptica with sequence similarities to plasmids from Gram-positive organisms. Mol Microbiol 6:1785–1799.

44. Newman JR, Fuqua C. 1999. Broad-host-range expression vectors that carry the L-arabinose-inducible Escherichia coli araBAD promoter and the araC regulator. Gene 227:197–203.

45. Bai J, Raustad N, Denoncourt J, van Opijnen T, Geisinger E. 2023. Genome-wide phage susceptibility analysis in Acinetobacter baumannii reveals capsule modulation strategies that determine phage infectivity. PLoS Pathog 19.

46. Landeta C, Mejia-Santana A. 2022. Union Is Strength: Target-Based and Whole-Cell High-Throughput Screens in Antibacterial Discovery. J Bacteriol. American Society for Microbiology 10.1128/jb.00477-21.

47. Heras B, Totsika M, Jarrott R, Shouldice SR, Gunčar G, Achard MES, Wells TJ, Argente MP, McEwan AG, Schembri MA. 2010. Structural and functional characterization of three DsbA paralogues from Salmonella enterica serovar typhimurium. Journal of Biological Chemistry 285:18423–18432.

48. Tinsley CR, Voulhoux R, Beretti JL, Tommassen J, Nassif X. 2004. Three homologues, including two membrane-bound proteins, of the disulfide oxidoreductase DsbA in Neisseria meningitidis: Effects on bacterial growth and biogenesis of functional type IV pili. Journal of Biological Chemistry 279:27078–27087.

49. Sinha S, Langford PR, Kroll JS. 2004. Functional diversity of three different DsbA proteins from Neisseria meningitidis. Microbiology (N Y) 150:2993–3000.

50. Raczko AM, Bujnicki JM, Pawłowski M, Godlewska R, Lewandowska M, Jagusztyn-Krynicka EK. 2005. Characterization of new DsbB-like thiol-oxidoreductases of Campylobacter jejuni and Helicobacter pylori and classification of the DsbB family based on phylogenomic, structural and functional criteria. Microbiology (N Y) 151:219–231.

51. Kpadeh ZZ, Jameson-Lee M, Yeh AJ, Chertihin O, Shumilin IA, Dey R, Day SR, Hoffman PS. 2013. Disulfide bond oxidoreductase DsbA2 of Legionella pneumophila exhibits protein disulfide isomerase activity. J Bacteriol 195:1825–1833.

52. Fenn K, Strandwitz P, Stewart EJ, Dimise E, Rubin S, Gurubacharya S, Clardy J, Lewis K. 2017. Quinones are growth factors for the human gut microbiota. Microbiome 5:161.

53. Kadokura H, Bader M, Tian H, Bardwell JC, Beckwith J. 2000. Roles of a conserved arginine residue of DsbB in linking protein disulfide-bond-formation pathway to the respiratory chain of Escherichia coli. Proc Natl Acad Sci U S A 97:10884–10889.

54. Thibodeau Stacey, Fang Rui, Joung J. Keith. 2004. High-throughput β-galactosidase assay for bacterial cell-based reporter systems. Biotechniques 36:410–415.

